# Phylogenetic and biogeographic history of the Snooks (Centropomidae: Carangiformes) spanning the closure of the Isthmus of Panama

**DOI:** 10.1101/2024.01.22.576679

**Authors:** Natalia Ossa-Hernández, Gustavo A. Ballen, Arturo Acero P., Jose Tavera

## Abstract

Amphiamerican New World fishes provide a unique opportunity to explore the impact of geological processes and the formation of geographic barriers on biological diversification across both spatial and temporal dimensions. We employed phylogenetic and biogeographic methods to assess the impact of the emergence of the isthmus of Panama on the evolutionary history of Snooks. Bayesian methods were used for phylogenetic inference and divergence time estimation, incorporating the fossil record of Carangidae, Centropomidae, Istiophoriformes, Latidae, and Sphyraenidae to establish a timeline using methods of stratigraphic intervals. Biogeographic models were explored to test the hypothesis that trans-Isthmian vicariant events are coeval with the Isthmus closure. Our results suggest a sister relationship between Centropomidae and Sphyraenidae with a common ancestor that originated in the Upper Cretaceous (∼78.18 Ma). The biogeographic model DEC+*j* indicated within-area speciation and dispersal (founder effect) as the primary modes of speciation in the genera *Centropomus*, *Lates,* and *Sphyraena*. The dispersion in the family Centropomidae was estimated from the Tropical Eastern Pacific to the Tropical Western Atlantic during the emergence of the isthmus of Panama. The alignment of divergence times with ancestral species distributions suggests a possible synchrony between the current distribution in *Centropomus* species and the gradual processes involved in the formation of the Isthmus of Panama during the Miocene. Furthermore, recent speciation events within each basin imply an influence of post-closure environmental conditions on the evolution of this group of fishes.

## Introduction

Geographic isolation plays a crucial role in speciation in different ways, such as reproductive isolation, adaptation to new environments, and other processes that either foster differentiation or restrict gene flow (Worsham et al. 2017). While the occurrence of speciation through geographic isolation is relatively rare in open systems like marine ecosystems, various ocean barriers exist, each exerting differing degrees of isolation. Soft barriers arise from oceanographic conditions or hydrobiological processes disrupting the interaction between conspecific individuals. Hard barriers are generally restrictions of terrestrial origin that physically separate marine populations (Cowman and Bellwood 2013), being the emergence of the Isthmus of Panama (IP) is one of its iconic examples. The rise of a terrestrial landscape separating the Eastern Pacific and the Western Atlantic led to the disappearance of the Central America Seaway (CAS), the creation of the Caribbean Sea, the modification of ocean dynamics, and therefore, an enormous impact on the Earth climate (Jaramillo 2018). This barrier acted as a bridge for terrestrial fauna and flora between South and North America and a barrier isolating the marine biota on both sides (Birkerland 1990; Duque-Caro 1990; Bartoli et al. 2005; Lessios 2008; Montes et al. 2012; Coates and Stallard 2013; Bacon et al. 2015; O’Dea et al. 2016).

Centropomidae *sensu stricto* is a monogeneric New World family of marine fish, consisting of 13 species all in the genus *Centropomus* Lacepède, 1802, that inhabit marine, estuarine, and mangrove ecosystems with sandy and rocky bottoms. Given its presence on both sides of the IP, the family Centropomidae provides an excellent opportunity to test the effects of a hard barrier on divergence times and biogeographic patterns. They are commonly known as snooks or *robalos* in various regions of Central and South America where they have great commercial importance (Vergara-Chen 2014). Snooks have a high tolerance to low salinity but are vulnerable to temperature extremes (Tringali et al. 1999; Anderson and Williford 2020). The morphology of centropomid species is conservative, with strong similarities among species. Jordan (1908) suggested three pairs of sister species of the family Centropomidae based on morphological similarities, all living under essentially the same environmental conditions but separated since the end of the Miocene by the rise of the IP.

Snooks belong to the series Carangaria, order Carangiformes, alongside the families Latidae and Sphyraenidae all of which share some morphological similarities (Betancur et al. 2017; Girard et al. 2020; Thacker and Near 2023). The phylogenetic relationship between these families is still a subject of debate. Centropomidae has been proposed to be sister to Latidae (Tringali et al. 1999; Li et al. 2011; Betancur-R. et al. 2013, 2017; Carvalho – Filho et al. 2019; Anderson and Williford 2020; Figueiredo-Filho et al. 2021), or Sphyraenidae (Near et al. 2013; Mirande 2017; Rabosky et al. 2018; Girard et al. 2020). Latidae, with three extant genera, is distributed in Indo-West Pacific and African freshwater basins, often exhibiting endemism to specific lakes (Otero et al. 2014). Sphyraenidae comprises 29 extant species known as barracudas, that are found in tropical and subtropical regions of the Atlantic, Indian, and Pacific Oceans (Fricke et al. 2023). Our understanding of the biogeographic patterns in this group of families remains unexplored. The influence of geographic barriers, such as the IP, on the ancestral distribution of these species is still uncertain. If the Isthmus closure played any role in speciation in this group, we would expect to find a common area with pre-closure ages and younger, post-closure distributions of ages. Here, we constructed a time-calibrated phylogeny focusing on Centropomidae, Latidae, and Sphyraenidae. We used probabilistic modeling of biogeographic events on the phylogeny to test the association of the closure of the IP in 1) the divergence time of the species and, 2) the ancestral distribution of the species to assess the role of the isolation and large-scale barriers in shaping the current species geographical distribution.

## Methods

### Sample collection

We collected specimens at local fish markets and landing areas across Caribbean and Pacific localities. All specimens collected were preserved and stored at the fish collection of Universidad del Valle (CIR-UV). Fishes were photographed, and a small fragment of the right pectoral fin was preserved in 96% ethanol and stored at -20 °C. We performed analyses using sequences generated in the present study and additional sequences obtained from GenBank. A total of 182 sequences (70 from this study and 112 from GenBank; Supplementary Table S1) from 13 species of Centropomidae, 22 species of Sphyraenidae, eight species of Latidae, two species of Carangidae, one species each of Coryphaenidae, Echeneidae, Nematistidae, Xiphiidae, and three species of Istiophoridae were used to estimate divergence times and infer phylogenetic relationships.

### DNA extraction, PCR amplification, and sequencing

We extracted DNA following the Salting Out protocol (Sambrook et al. 1989). Four mitochondrial genes (12SrRNA, 16SrRNA, Cytochrome oxidase subunit I – COI, and cytochrome b -CYB) and one nuclear marker, a single-copy locus TMO-4c4 were amplified. The PCR amplification reactions were conducted in a final volume of 15µl containing 1 µl of DNA of stock concentration, 1.5 µl 10X reaction buffer BD (0.8M Tris -HCL, 0.2 M(NH_4_)_2_SO_4_; Solis Bio Dyne), 1.2 µl of 2.0 mM MgCl_2_, 0.4 mM premixed deoxynucleotide triphosphates, 0.1 µl of 5Uml *Taq* FIREPOL DNA polymerase (Solis Bio Dyne), and 0.2 µl of each oligonucleotide primer, each at 20mM concentration. PCR cycle parameters used to amplify all the genes included an initial denaturation step at 94 °C for 3min, followed by 35 cycles at 94 °C for 45 s, 48 °C – 60°C (see supplementary Table S2 for details of the primers sequences and annealing temperature used for each gene) for 45 s, and extension at 72 °C for 45 s, and a final extension step at 72°C for 10min. All PCR products were loaded and run in agarose gels at 1% to verify the correct amplification. Macrogen Inc. (Seoul, Korea) performed the standard sequencing service.

### Phylogenetics analysis: Bayesian inference

Sequences were edited using Geneious (Kearse et al. 2012) and aligned using MAFFT algorithm (Katoh et al. 2013). The matrix concatenation was carried out using phyx (Brown et al. 2017). The substitution model for each gene was selected based on the Akaike information criterion corrected for small sample sizes (AICc) using jModelTest 2 (Darriba et al. 2012). The substitution models that best fitted each locus were GTR + I +G (12SrRNA, 16SrRNA, COI, CYB), and HKY+I + G (TMO4c4).

Parallel-tempering Metropolis-Hastings MCMC (Metropolis et al. 1953; Hastings 1970) was used for sampling from the posterior distributions of the phylogenetic model. Sampling was carried out in Mr. Bayes (Ronquist et al. 2012) using parallelism (Altekar et al. 2004). We ran four independent analyses, each with eight chains (one cold and seven hot) for 40,000,000 generations, sampling every 4,000th generation. A summary of posterior distributions was carried out after applying a burn-in of 50% while combining the remainder of the four analyses. Inter-chain parameter convergence was assessed using the potential scale reduction factor which approached 1.0 for all parameters. Intra-chain parameter convergence was assessed using the effective sample size, which was >1,000 for all parameters. The average standard deviation of split frequencies was used for assessing topological convergence, which reached a value of 0.02 at the end of the analysis. The posterior tree distribution was summarized using a majority-rule consensus tree with average branch lengths. Branch support values are posterior probabilities summarized from the posterior tree distribution.

### Fossil calibrations and stratigraphic intervals

Node calibrations based on the fossil record are often problematic because they represent a single constraint on the age of the node, which results in an improper distribution [e.g., Uniform (minimum age, Infinite)] which does not integrate to 1.0 and often leads to convergence issues (Yang 2014). However, it is also difficult (and sometimes impossible) to set a maximum that does not depend for instance on a general constraint (e.g., the root age) that will define all the node calibrations in the tree. This issue does not apply only to hard bounds: It is also impossible to define a distribution with soft bounds unless we set at least two quantiles so that we can fit parameters that define a distribution matching these values. When we use a single constraint, we have only a single quantile and no proper distribution.

It is possible to estimate both the origination and extinction times for a given lineage using stratigraphic intervals, which are models that describe these times as parameters that are a function of the pattern of fossil occurrences in time (Strauss and Sadler 1989). Multiple methods exist and are based on different assumptions about the pattern or process that generates the fossil occurrences (Marshall 2010). One of these methods is the Beta-adaptive model (Wang et al. 2016) which can be used when the preservation potential is unknown given a sample of occurrences in time for a given lineage. The advantage of these methods is that we can define a proper statistical distribution for the origination time of a series of occurrences, and therefore use the known fossil record of a given lineage as a sample for estimating the origination parameter as a distribution. Here, we are equating the origination time of a lineage with its most recent common ancestor. Thus, we can specify node time distributions using the fossil record, regardless of its completeness.

We used the Beta-adaptive method and the known fossil record of the families Carangidae (35 occurrences), Centropomidae (5 occurrences), Istiophoridae (52 occurrences), Latidae (11 occurrences), and Sphyraenidae (76 occurrences) (Fig. 1). A survey of published records was used to construct the sample of time occurrences for the families Carangidae, Istiophoridae, and Latidae (Uhen et al. 2023; website: https://paleobiodb.org/). Time occurrences of the family Sphyraenidae are based on Ballen (2020) and direct examination of museum specimens. The time occurrences of the family Centropomidae are herein published for the first time from a review of the literature as well as new fossil occurrences (Supplementary material).

All the information associated with these records can be found on the website https://www.floridamuseum.ufl.edu/vertpaleo-search/. The resulting distribution for the origination time of each family was used to generate the posterior credible interval, which in turn was approximated by the quantiles 0.025 and 0.975 of a truncated Cauchy distribution (Inoue et al. 2010) which is implemented for defining node calibrations in MCMCTree.

**Figure 1.**
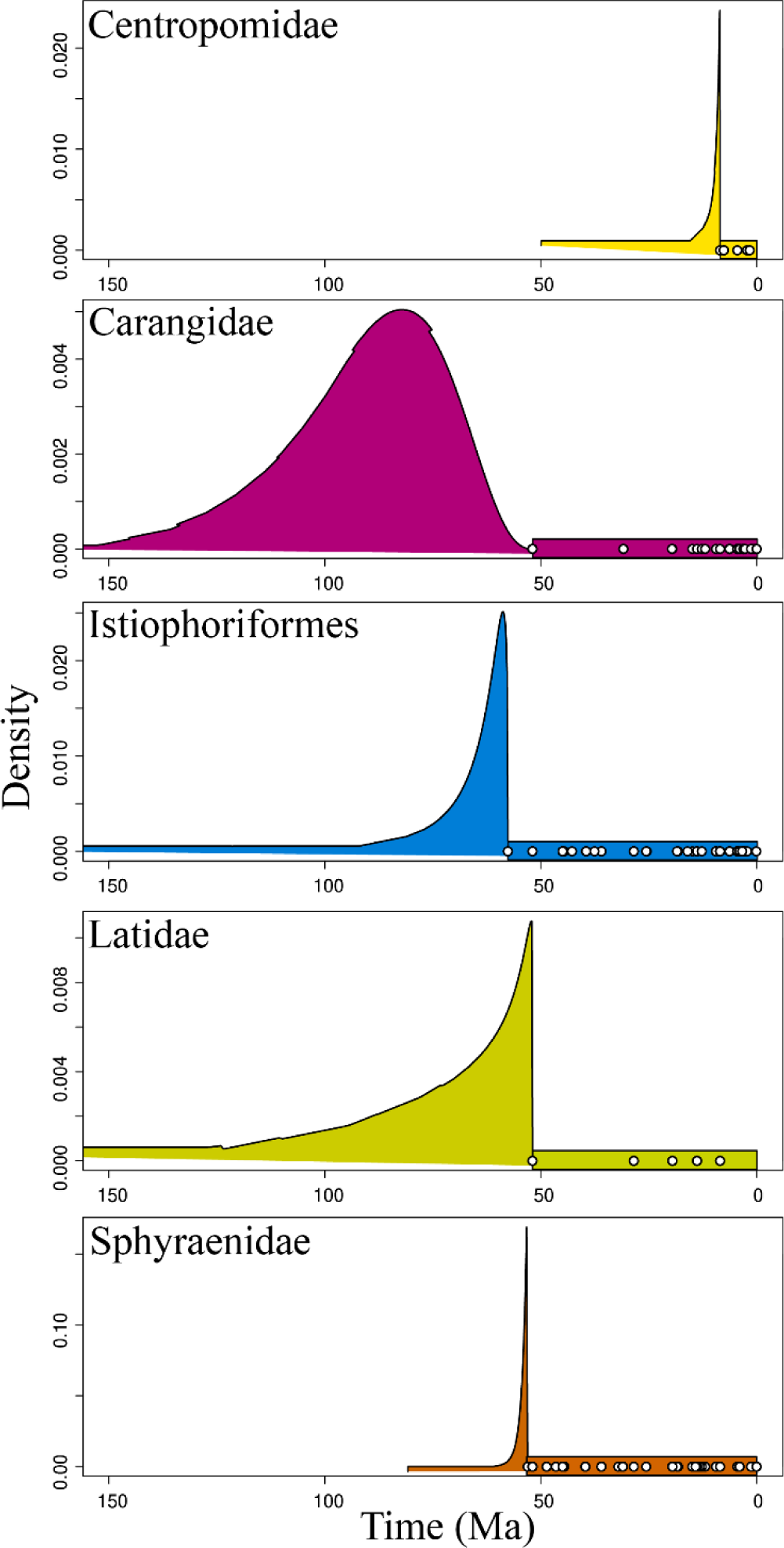
Stratigraphic series for the families Centropomidae, Carangidae, Latidae, Sphyraenidae and some families in the order Istiophoriformes. Circles represent fossil occurrences in geologic time, and shaded areas represent the posterior density on the origination age of each stratigraphic series. Note that all the series are constrained to be extant, and therefore theta2 is fixed to 0.0 (present time). The extent of the posterior on origination time depends on the number of observations as well as on the temporal pattern of occurrences: If they become less frequent towards the origination time, the density has higher variance, on the other hand, if occurrences are more frequent towards the origination time, the posterior has lower variance. The vertical axis (density) is not scaled across panels, whereas the horizontal (time) axis is shared across panels. The posterior densities on the origination time theta1 are used as node calibration densities in the divergence time estimation analysis.

### Divergence time estimation

We used divergence time estimation to put the phylogenetic hypothesis in a temporal context. We used a summary maximum credibility tree, that is, the topology with the maximum sum of branch posterior probabilities as the target tree. We calculated that tree with logcombiner (Bouckaert et al. 2019). The analysis was carried out in MCMCTree of the PAML suite (Yang 2007). We used the method of approximate likelihood of dos Reis and Yang (2011) under the HKY substitution model to speed up the analysis runtime.

The clock model is important for the estimation of node ages (dos Reis et al. 2018), and therefore we used Bayesian model selection via marginal likelihood calculation using stepping stones for choosing the best relaxed-clock model for this dataset. We used the stepping stones procedure implemented in the mcmc3r package (dos Reis et al. 2018) R package (R Core Team 2021) and the block bootstrap calculation of the standard error proposed by Álvarez-Carretero et al. (2022).

We used two clock models (independent and autocorrelated), and 64 stones. Each stone was sampled with a burn-in of 4,000 generations, and then sampling every two until 10,000 samples were reached. The large number of stones compensates for the shorter chains when marginalizing the likelihood over the model parameters. The relaxed clock model was set to independent rates from a lognormal distribution following the results of the model selection analysis (Model Posterior Probability = 1.0; Supplementary Table S3).

We used a birth-death tree model setting the birth and death rates to 1.0, and the sampling fraction to 0.1. We used a gamma-Dirichlet (2,20) prior to the rates and the gamma-Dirichlet (1,10) prior to the sigma^2^ parameter of the relaxed clock model. We set a gamma (2,0.2) prior to the kappa and a gamma (2,4) prior to the alpha parameters of the substitution model.

Five node calibrations and an age constraint on the root were used to generate the time prior: The nodes corresponding to the most recent common ancestor of the families Carangidae, Centropomidae, Latidae, and Sphyraenidae, and the order Istiophoriformes. Node calibrations were specified as truncated Cauchy distributions as described in the previous section. A soft-bound maximum on the root age was set to 120 Ma. Four independent analyses were run and compared to assess across-chain convergence. Effective sample size (ESS) values larger than 500 were found and suggest adequate mixing inside each chain. All the analyses were run on the *Brycon* server at IBB/UNESP Botucatu.

### Biogeographic modeling and ancestral range estimation

Ancestral areas of the families Centropomidae, Latidae, and Sphyraenidae were estimated on the time-calibrated phylogeny using BioGeoBEARS (Matzke 2013). Three models were tested: Dispersal-Extinction-Cladogenesis (DEC, Ree and Smith 2008), a likelihood version of the parsimony-based Dispersal Vicariance analysis (DIVALIKE, Ronquist 1997), and a likelihood version of the range evolution model implemented in BayArea (BAYAreaLIKE, Landis et al. 2013). In each model, an additional “j” parameter (founder event/jump speciation) was added, which allows descendant lineages to have a different area from the direct ancestor (Matzke 2013). Thus, a total of six models were tested and compared using statistical fit with the corrected Akaike Information Criterion (AICc). Species distribution data were obtained from Robertson and Allen (2015), Froese and Pauly (2017), and Robertson and Van Tassell (2023). Six marine bioregions were used based on Kocsis et al. (2018): Tr. EP: Tropical Eastern Pacific, Tr. WA: Tropical Western Atlantic, Tr. EA: Tropical Eastern Atlantic, Tr. IP: Tropical Indo-Pacific, Af: African; Au: Temperate Australian. Thus, we set six as the maximum number of areas in the ancestral area reconstruction analyses.

### Biogeographic stochastic mapping

We estimated the number of each type of biogeographic event in the phylogeny using the Biogeographical Stochastic Mapping (BSM) implemented in BioGeoBEARS (Matzke 2015; Dupin et al. 2016). Six types of biogeographic events possible under the models, within-area speciation (sympatry), vicariance, and dispersal events (founder events) were tested. We conducted BSM using the MCC tree and the DEC + *j* that produced a significantly better fit to the data compared with other tested models. We generated 50 stochastic maps, and the event frequencies were estimated by taking the mean and standard deviation of event counts from 50 BSM.

### Code and data

All the code and data necessary for reproducing the results are available on Zenodo (DOI https://doi.org/10.5281/zenodo.10535162) as well as on https://github.com/gaballench/centropomidae_divtime.

## RESULTS

### New fossil occurrences of the Centropomidae

We reviewed the fossil specimens of the family Centropomidae deposited at the Florida Museum of Natural History (FM). A total of 26 bones were analyzed through the pictures processed by the museum (Supplementary material). The photographs from FM were compared with literature referent at the family Centropomidae (Fraser 1968; Potthoff and Tellock 1993) and with the same bones in extant species. The oldest record dates to the upper Late Miocene (5.33 - 11.63 Ma) and the earliest to Pleistocene (Calabrian 0.774 – 1.80 Ma). Fossils were collected in the Hawthorn Formation (15 occurrences) in Polk, Florida, the Tamiami Formation (three occurrences) in Southwest Florida, the Alachua Formation (four occurrences) north of Newberry, western Alachua County, and the Bermont Formation (four occurrences) Hillsborough County, Florida. The material corresponds to the head and mandible bones mainly.

### Phylogenetic inference

In the concatenated analysis, a total of 3234 base pairs (bp) from combined genes (12SrRNA – 832 bp, 16SrRNA – 569 bp, COI – 614 bp, CYTB– 782 bp, TMO4c4 – 437 bp) were used to reconstruct the phylogeny of three families: Centropomidae (snooks), Latidae (perches), and Sphyraenidae (barracudas). Our findings indicate the monophyly of each family, supported by a posterior probability of 1.0. Centropomidae and Sphyraenidae were recovered as sister clades with a posterior probability 0.91, while Latidae was recovered near the base of the tree (Fig. 2).

### Divergence time estimation

The estimated crown age for the family was 58.47 Ma (95% highest posterior density interval, HPDi= 50.65 – 72.69). The genus *Lates* displayed an estimated divergence age of 26.23 Ma (95% HPDi= 17.27 – 34.77) while the genera *Psammoperca* and *Hypopterus* exhibited an estimated divergence age of 26.83 Ma (95% HPDi= 6.34 – 55.9). Two pairs of sister species were recovered: *L. calcarifer – L. japonicus* with a divergence time of 22.16 Ma (95% HPDi= 13.43 – 31.02) and *L. angustifrons – L. microlepis* with a more recent divergence time of 310 Ka (95% HPDi= 0 – 0.92).

The estimated crown age for the family Centropomidae was 32.91 Ma (95% HPDi= 24.59 – 40.47). Two main clades encompassing the 13 species of the family were identified. The first one with a high support (0.99 PP) and an age of 26.69 Ma (95% HPDi= 24.59 – 40.47) comprised ((*C. medius*, *C. pectinatus*), (*C*. *mexicanus*, *C. parallelus* (*C. nigrescens*, (*C. viridis*, (*C. poeyi* (*C. irae*, *C. undecimalis*)))))). Noteworthy in this clade are three sister species pairs: a transisthmian pair (*C. pectinatus* – *C. medius*) with a divergence age estimated of 6.2 Ma (95% HPDi= 2.95 -9.74), a sympatric pair found in the Western Atlantic (WA) (*C*. *mexicanus*, *C. parallelus*) with a recent divergence time (estimate= 1.18 Ma, 95% HPDi=0.27 – 2.26), and an allopatric pair, *C. irae* restricted to Amapa, Brazil, and *C. undecimalis* with broad distribution in the WA with a divergence time estimated at 5.51 Ma (95% HPDi=2.97 – 8.48). The second clade supported by one PP includes the species (*C. ensiferus,* (*C. unionensis,* (*C. robalito,* and *C. armatus*))). Within this clade, the sister species *C. armatus* and *C. robalito* are both present in the EP, and represent a sympatric pair with the most recent time of divergence, estimated age of 210 Ka (95% HPDi=0 – 0.51).

The crown age for the family Sphyraenidae was estimated to be 58.39 Ma (95% HPDi= 58.34 – 58.36). Two main clades can be identified. The first one was supported by a posterior probability of 0.92 with an age of 53.47 Ma (95% HPDi= 48.9 – 57.18) which included the species ((*S. obtusata, S. flavicauda*)*, (S. pinguis,* (*S. iburiensis, S. chrysotaenia*), (*S. idiastes,* (*S. viridensis, S. argentea*), (*S. helleri,* (*S. novaehollandiae,* (*S. africana,* (*S. japonica,* (*S. acutippinis, S. waitii*)))))))))). The second one is supported by a posterior probability of 1.0 and has an age of 31.31 Ma (95% HPDi= 22.26 – 40.92) including the species (*S. putname,* (*S. qenie,* (*S. barracuda, S. jello*), (*S. forsteri,* (*S. ensis,* (*S. borealis, S. guachancho*)))))). Six sister-species pairs were identified. The divergence time in sister pairs ranges between 1.2 to 9.66 Ma. *S. guachancho* – *S. borealis* (estimate= 1.2, 95% HPDi=0.22 – 2.37); *S. waitii – S. acutipinnis* (estimate= 6.98, 95% HPDi=1.68 – 12.99); *S. jello – S. barracuda* (estimate= 7.23, 95% HPDi=2.45 – 12.72); *S. argentea – S. viridensis* (estimate= 8.5, 95% HPDi=1.34 – 18.05); *S. chrysotaenia – S. iburiensis* (estimate= 9.66, 95% HPDi=1.18 – 18.97) and, *S. flavicauda* – *S. obtusata* (estimate= 10.37, 95% HPDi=2.56 – 19.52).

**Figure 2.**
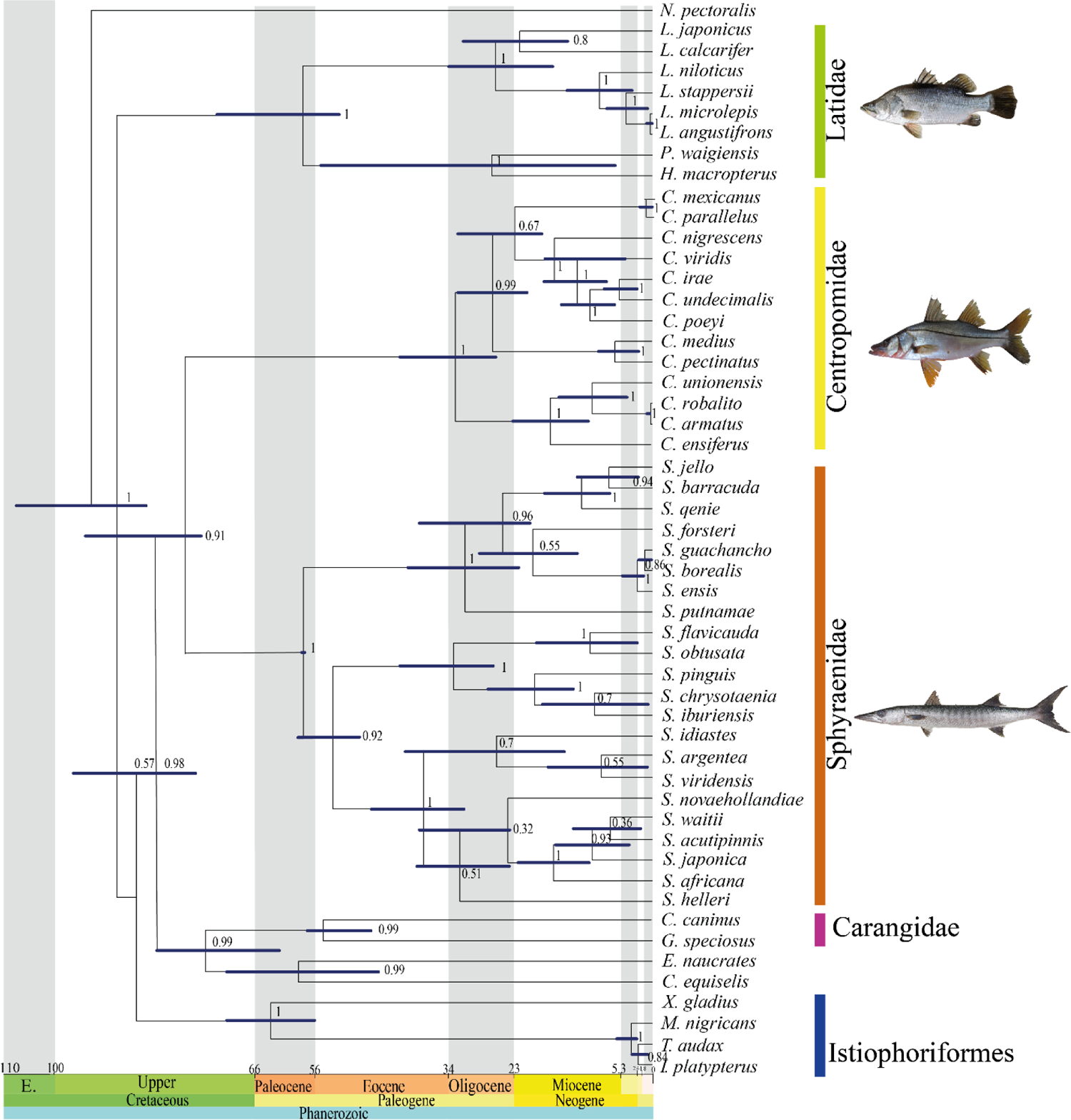
Time-calibrated maximum credibility tree of the family Centropomidae, Latidae, and Sphyraenidae. Blue bars represent 95% highest posterior density interval (HPDi) of the node ages. Node labels represent the posterior probability.

### Biogeographic analysis

Historical range optimization using BioGeoBEARS indicates that the biogeographic model DEC+*j* was the best one, based on the Akaike information criterion (weight 1.0; Supplementary Table S4). The founder event speciation parameter (*j*) was favored in all models tested, indicating that this mode of dispersal was significant in forming the broad-scale biogeographic pattern in these families (Supplementary Table S4).

The time-calibrated phylogeny suggests that the common ancestor of the clade Centropomidae + Sphyraenidae arose during the upper Cretaceous from approximately 88.49 Ma to 69.01 Ma (Fig. 3). Ancestral range estimations under this best-fitting model (DEC*+j*) showed that the most probable ancestral area for extant species of Sphyraenidae + Centropomidae was the Tr. IP P =0.25, (with P= 0.26 for Tr. EP and P= 0.21 for the combination Tr. EP + Tr. WA) (Fig. 3). The node containing all species of the family Sphyraenidae has the Tr. IP (P=0.57) as the more probable estimated ancestral range. Other deep nodes in the family Sphyraenidae also have Tr. IP as the estimated ancestral range. During the Oligocene (26 Ma) the estimated ancestral area showed a clade in the Tr. IP + Tr. EP (P=0.66) and another one in the Tr. EP (P=0.92). Ancestral areas in the family Centropomidae have the Tr. EP + Tr. WA as the estimated range (P= 0.58) (Fig. 3). Two clades with three species each, diverged during the Miocene (10 Ma) with the first having an estimated ancestral area in the Tr. EP (P=0.97) and the other in the Tr. WA (P= 0.98). The root node of the family Latidae has an ancestral area in the Tropical Indo Pacific during the Paleocene (58.47 Ma), and during the Miocene one clade has an estimated ancestral in the freshwaters of Africa (*P=*0.97).

**Figure 3.**
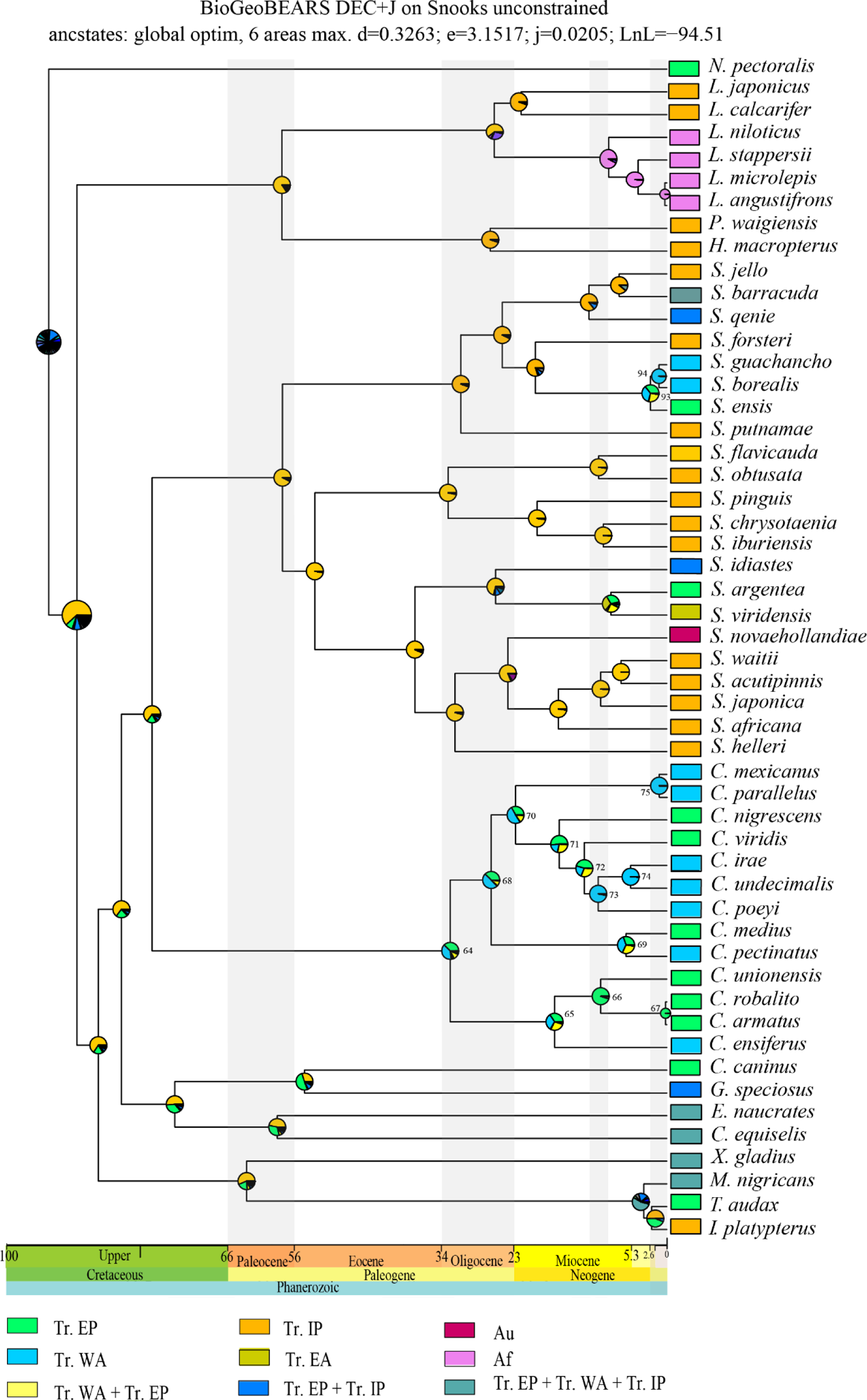
Biogeographic history based on the BioGeoBEARS optimization of the time-calibrated phylogeny. Colored boxes at the terminals of the phylogeny show the extant geographic distribution of species. Operational areas, according to Kocsis et al. (2018) are Tr. IP: Tropical Indo-Pacific; Tr. WA: Tropical Western Atlantic; Te. Au: Temperate Australian; Tr. EP: Tropical Eastern Pacific; Tr. EA: Tropical East Atlantic; Af: African. Other biogeographic areas are based on combinations of those defined a priori. The numbered nodes are the ones with estimated ancestral area in Tr. EP, Tr. WA, and Tr. EP or Tr. WA. Shaded areas represent periods of major divergence events of target families.

### Estimation of the number, type, and directionality of biogeographical events

A summary of our Biogeographical Stochastic Models (BSMs) revealed that most biogeographical events across the species included in the present study were within–area speciation (58%), followed by dispersal (38%), and a few vicariant events (4%) (Table 1). The largest number of events of within–area speciation occurred 53% in Tr. IP, 21 in % Tr. EP, and 16% in the Tr. WA. Twelve nodes were examined in detail in the family Centropomidae finding that 68% of the events correspond to within-area speciation, 24% to dispersal, and 8% to vicariance. Dispersal events were estimated at 60% from the Tr. EP to Tr. WA during the Miocene. Twenty-one nodes were examined in the family Sphyraenidae finding that 85% of the events correspond to within-area speciation, 13% to dispersal, and 2% to vicariance. Seven nodes in the Latidae clade showed that 85% of the events correspond to within-area speciation, 13% to dispersal, and 2% to vicariance (Table 1).

**Table 1.** Summary of biogeographical stochastic mapping counts using the DEC+ *j* model. Mean numbers of the different types of events estimated are shown here along with standard deviations. No range contractions were estimated because the relevant model parameter (e) was not required in the best-fitting model.

| Mode | Type | Total mean<br>(SD) | % | Centropomidae<br>Mean (SD) | % | Sphyraenidae<br>Mean (SD) | % | Latidae<br>Mean (SD) | % |
| --- | --- | --- | --- | --- | --- | --- | --- | --- | --- |
| **Within-area speciation** | Speciation | 30.4<br>(2.15) | 47 | 40.4<br>(13.13) | 61 | 40.4<br>(12.53) | 72 | 48.2<br>(4.49) | 83 |
|  | Speciation subset | 7.4<br>(2.83) | 11 | 10.8<br>(2.22) | 7 | 13.3<br>(7.50) | 13 | 6 | 2 |
| **Dispersal** | Founder events | 10.3<br>(1.79) | 38 | 20.9<br>(16.43) | 24 | 12.8<br>(17.95) | 13 | 18.5<br>(21.92) | 10 |
| **Vicariance** | Vicariance | 2.8<br>(1.56) | 4 | 9.4<br>(5.73) | 8 | 2.7<br>(1.97) | 2 | 9<br>(9.90) | 5 |
| **Total** |  | 65.5<br>(1.56) | 100 | 150.0<br>(150.41) | 100 | 237.5<br>(303.32) | 100 | 87.5<br>(134.94) | 100 |

### Summary of estimated ancestral area over time in the Tropical Eastern Pacific and Tropical Western Atlantic during the Neogene

A total the 14 nodes in the phylogenetic inferences showed estimated ancestral areas in Tr. EP (2), Tr. WA (4), and Tr. EP or Tr. WA (8) as the estimated ancestral range during the Neogene. We extracted the age of each node with distribution in these areas and plotted this data. We observed that the distribution of areas is aligned with the final gradual process of emergence of the Isthmus of Panama where the Pacific and the Atlantic Oceans were separated. The nodes number 64 – 75 correspond to the family Centropomidae, where those nodes had an ancestral area in the Tr. EP + Tr. WA, Tr. EP, and Tr. WA with divergence times in the Oligocene (33.9 Ma – 23.03 Ma) to the Upper Miocene (11.63 Ma – 5.33 Ma) (Fig. 4), and the nodes with an estimated distribution either in the Tr. EP or in the Tr. WA has ages ranging between the Pliocene (5.33 Ma-2.58 Ma) to the Pleistocene (2.58 Ma-0.774 Ma) (Fig. 4). The nodes number 93-94 correspond to the family Sphyraenidae, where there is one node with ancestral areas in Tr. EP or Tr. WA with divergence during the early Pleistocene (∼2.49 Ma).

**Figure 4.**
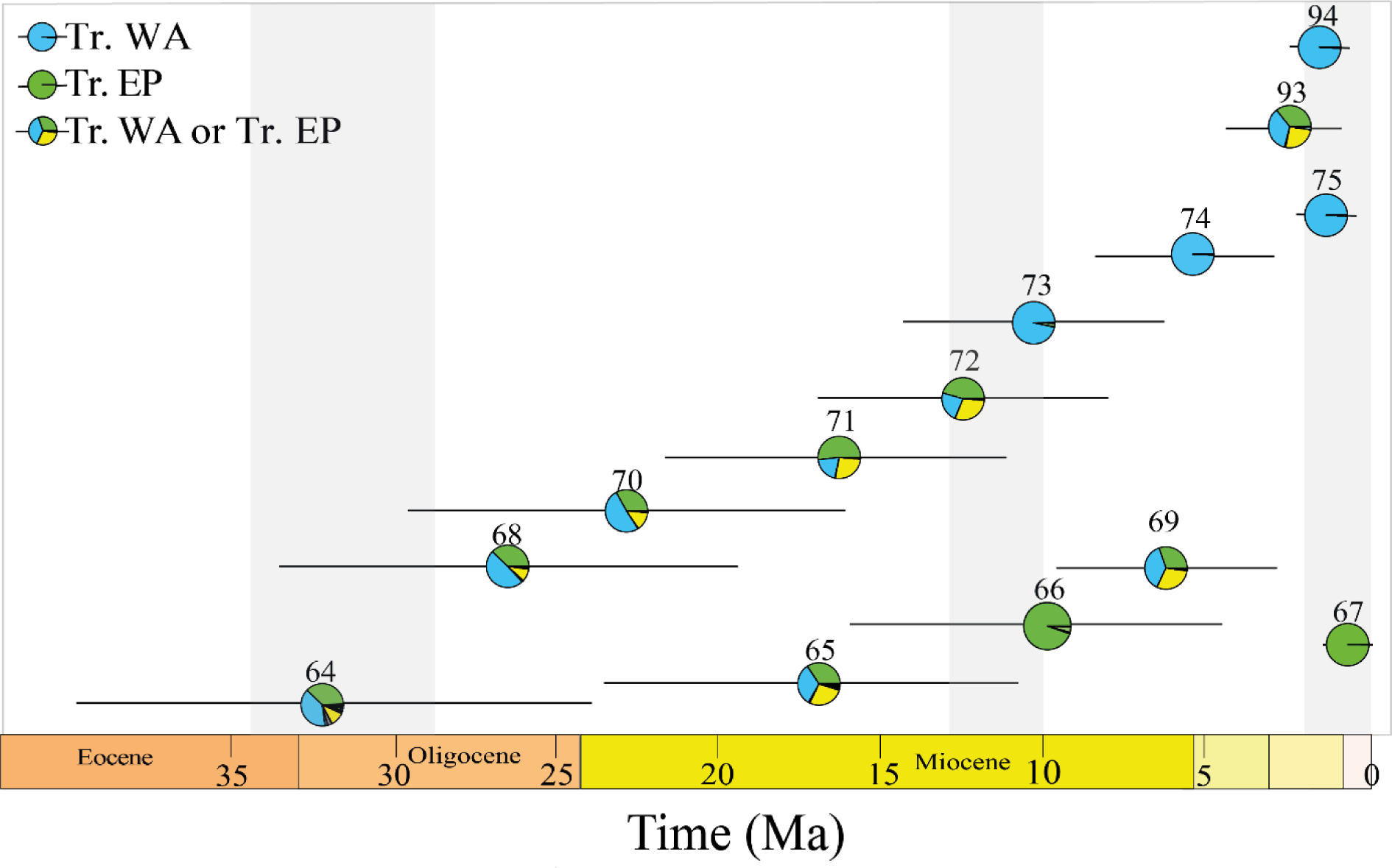
Forest-plot of value to High Posterior Density interval (HPDi) of the age of the 14 nodes in the phylogenetic inferences with estimated ancestral area in Tr. EP, Tr. WA, and Tr. EP or Tr. WA. Pie represents the estimated median age, and the length of the line represents the 95% HPDi. Gray shading as in Figure 3.

## Discussion

### Phylogenetic interrelationships between Centropomidae, Latidae, and Sphyraenidae

The historical debate and subsequent modifications regarding the interrelationship of the Centropomidae with the family Latidae have a long history. At the beginning of phylogenetic studies, Centropomidae comprised two subfamilies, Centropominae and Latinae (Greenwood 1976). A taxonomic and phylogenetic revision redefined the family Latidae as monophyletic, encompassing the genera *Lates, Psammoperca,* and *Eolates* (fossil), while the genus *Centropomus* was assigned to the family Centropomidae (Otero 2004).

Subsequently, the monophyly of the Centropomidae and Latidae clade was proposed by Li et al. (2011), although with weak support (0.74 PP), and only four species representing these two families were used. Betancur et al. (2013, 2017) recovered Latidae as the sister clade of Centropomidae with weak support (0.61 PP) and placed them as *incertae sedis* at the ordinal level as a part of Carangaria together with the family Sphyraenidae. Girard et al. (2020) recovered Centropomidae as a sister clade of Sphyraenidae and included them together with Latidae in the order Carangiformes.

After the designation of Centropomidae *sensu stricto*, little attention was directed to the interrelationships among these families. Interestingly, the family Sphyraenidae is related to the family Centropomidae in some studies (Near et al. 2013; Mirande 2017; Rabosky et al. 2018; Girard et al. 2020) (Fig. 5-A). We found that Centropomidae and Sphyraenidae are sister clades (0.91PP), while Latidae emerged as a basal unrelated clade, sister to all families included in our phylogeny. The phylogenetic hypotheses previously presented for the family Centropomidae had suggested Latidae as a sister family (Tringali et al. 1999; de Oliviera et al. 2014; Carvalho-Filho et al. 2019). In the same way, in a phylogenetic hypothesis of the Latidae, Centropomidae was used as an outgroup (Koblmuller et al. 2021). However, Santini et al. (2015) used Centropomidae as an outgroup in the Sphyraenidae phylogeny, being congruent with our results.

**Figure 5.**
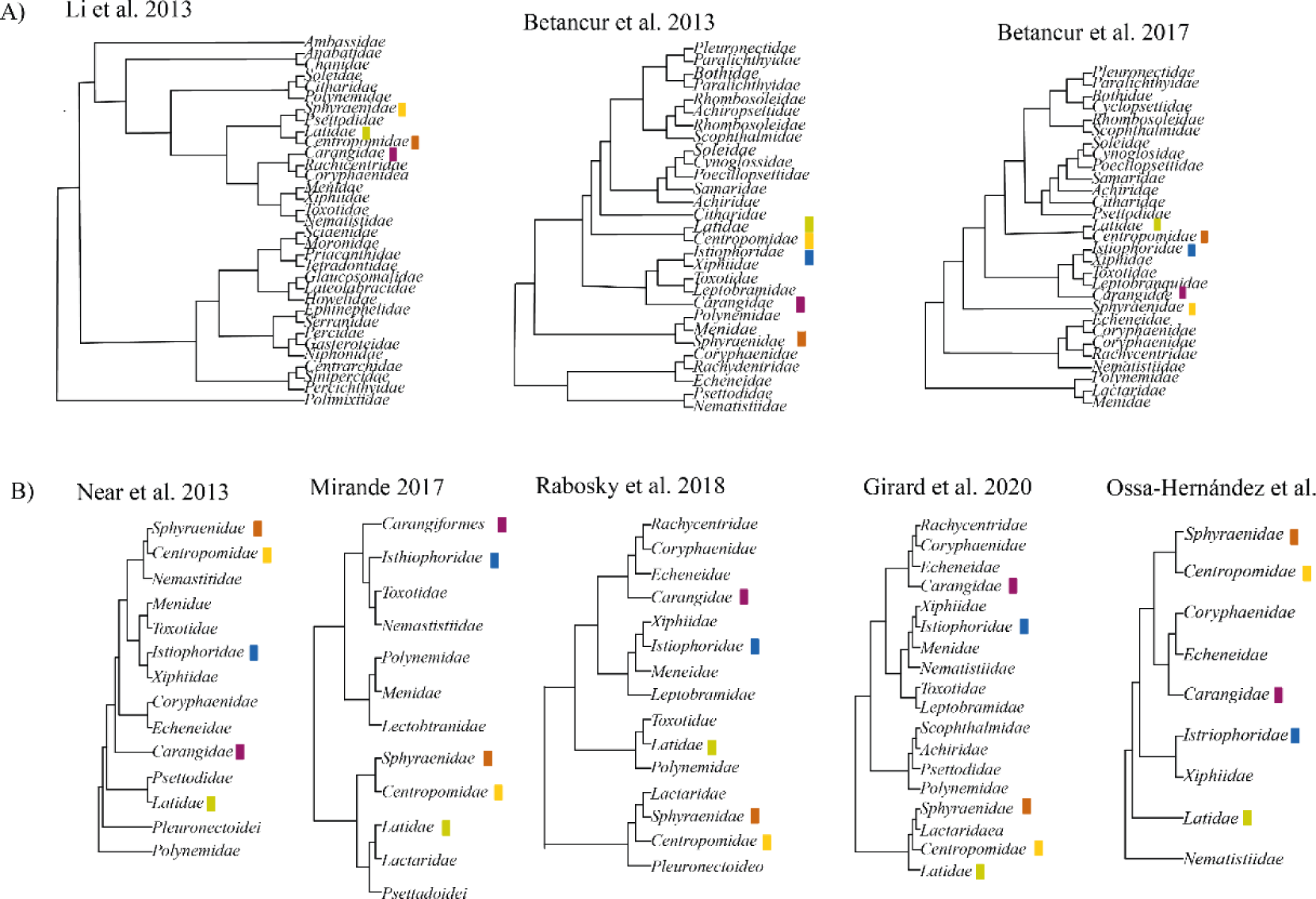
Hypotheses of relationships among ray-fin fishes where the closest relationship A) Centropomidae and Latidae are sister clade according to molecular studies (Li et al. 2011; Betancur 2013, 2017). B) Centropomidae and Sphyraenidae are recovered as sister clades based on the following molecular studies: Near et al. (2013); Mirande (2017); Rabosky et al. (2018); Girard et al. (2020). The color boxes indicate the families used in the present study.

### Hypotheses of the relationship among centropomid species

This study represents the first attempt at estimating the divergence times among species of the Centropomidae by using information derived from the fossil record. De Oliviera et al. (2014) recovered an age for the Centropomidae family during the Miocene (∼23 Ma), whereas our findings place the family’s origin in the Oligocene (∼32.91 million years ago). Additionally, the divergence times for sister species pairs do not align with De Oliviera et al. (2014).

We have identified two main clades congruent with the findings of Tringali et al. (1999); however, there are discrepancies in the relationships within each clade. It is important to note that Tringali et al. (1999) based their phylogenetic inferences on one locus. In the first clade ((*C. medius, C. pectinatus*), (*C. mexicanus, C. parallelus* (*C. nigrescens*, (*C. viridis*, (*C. poeyi* (*C. irae, C. undecimalis*)))))), our topologies align with Tringali et al. (1999). Nonetheless, we include *C. irae,* a species described after Tringali and collaborators published their hypothesis. In our phylogeny, this species is retrieved as a sister to *C. undecimalis*—a sister pair also identified by Oliveira et al. (2014). In the second clade (*C. ensiferus,* (*C. unionensis,* (*C. robalito,* and *C. armatus*))), we report *C. armatus* and *C. robalito* with PP 1.0. While Tringali et al. (1999) recovered *C. robalito* and *C. ensiferus* with 78% bootstrap support and *C. armatus* and *C. unionensis* with 98% bootstrap. Oliveira et al. (2014) also identified *C. robalito* as a sister species to *C. ensiferus*. Figueiredo-Filho et al. (2021) suggest that *C. robalito* and *C. armatus* are the same species based on the gene COI; however, our findings based on five molecular markers suggest that both are distinct species, yet with a very recent divergence time. Figueiredo-Filho et al. (2021) carried out a phylogenetic and taxonomic revision of the genus *Centropomus* with an emphasis on Atlantic species. Their inferences based on COI led them to propose taxonomic modifications that do not align with our results. For example, they suggest *C. mexicanus* is a junior synonym of *C. parallelus;* however, our findings indicate that both are distinct species with a recent divergence time, a hypothesis supported also by Seyoum et al. (2023), which indeed proposed the existence of three lineages in the *C. mexicanus* and *C. parallelus* complex. Figueiredo-Filho et al. (2021) suggest that *Centropomus nigrescens* and *C. viridis* may be the same species, but our results indicate that these two species are distinct, lacking a close phylogenetic relationship and possessing different ages—*C. viridis* ∼12.54 Ma and *C. nigrescens* ∼16.35 Ma. Results are supported by Martínez-Brown et al. (2021), who conducted a morphological review, identifying diagnostic characteristics that differentiate these two species as distinct entities.

### Hypotheses of the relationship between sphyraenid species

The relationship between the different species of the family Sphyraenidae has been published previously. Santini et al. (2015) presented a phylogenetic hypothesis based on two mitochondrial genes (COI, CytB) of the family recovering three main groups. The same clades are recovered in our hypothesis (using five molecular markers), with some different relationships between species. The inclusion of *S. qenie* and *S. waitii* modified the interrelationship between the *S. barracuda* group and the *S. sphyraena* group reported by Santini et al. (2005). We recovered *S. jello -S. barracuda* as a sister species, this pair was previously reported by Betancur et al. (2017) in their ray-finned fish phylogenetic study.

### Ancestral range estimations

Our estimation of biogeographic history showed a similar pattern to the current distribution of the extant species. The families Latidae and Sphyraenidae are mostly distributed in the Eastern Hemisphere; species of the family Latidae are distributed in the Indo-Pacific and Africa; while species of the family Sphyraenidae are mainly distributed in the Indo-Pacific, with some species distributed in the Eastern Pacific and Western Atlantic and one species in Australia. On the other hand, in the Western Hemisphere, we have the family Centropomidae which is a New World endemic family.

Tropical IP was estimated as the ancestral area of the family Latidae and Sphyraenidae during the Paleocene, approximately ∼ 58 million years ago. However, when these families originated, the configuration of the Indo-Pacific differed from its present state. At that time, the Tethys Sea (TS) occupied the region currently comprised of the Mediterranean Sea and the Indo-Pacific Ocean. The TS exerted a substantial influence on the Earth’s ecological dynamics and supported a diverse array of marine and freshwater species (Hou and Li 2018, Zhao et al. 2022). The Indian Ocean had taken on its present configuration since 36 million years ago. During the Eocene–Oligocene boundary (∼33.9 Ma), when the southern Mediterranean was created, the connection between the TS and the Indian Ocean was reduced (Hou and Li 2018), until the complete closing of the Tethys during the Miocene (∼20). The closure of the TS during the Miocene (∼20 million years ago) was coeval with significant events of divergence in Latidae and Sphyraenidae. During that period the estimated ancestral area for the genus *Lates* was Africa, and for some species of the genus *Sphyraena,* it was the Tr. EP, Tr. EA, and Tr. WA. More recent divergence events were estimated in Sphyraenidae (∼2.49 Ma) in the Tr. EP + Tr. WA, this divergence occurred after the interruption of water exchange between the Pacific and Atlantic Oceans, following the complete formation of the Isthmus of Panama (∼3.5 Ma).

In the Western Hemisphere, the emergence of the Isthmus of Panama stands out as a pivotal geological event. Throughout the Oligocene to the Miocene, spanning from 33.9 to 15.98 million years ago, North and South America remained separated by one or more extensive and deep seaways that connected the Atlantic and Pacific Oceans (Jaramillo 2018). The snook family most probably originated in the Oligocene (∼32.91) in waters that divided North and South America in the absence of any geographical barriers. The events of diversification during the Oligocene to the early Miocene in the family Centropomidae overlap with terrestrial landscape development in the IP (Montes et al. 2012, Coates and Stallard 2013, Jaramillo 2018). The nodes with estimated ancestral area in Tr. EP or Tr. WA occurred in a range between 22.99 Ma to 12.54 Ma. The events of divergence with species restricted to either the Pacific or the Caribbean occurred ∼10 Ma. During this time (∼10 Ma), the closure of the CAS significantly affected the oceanic exchange between the Pacific and the Caribbean, decreasing the flow of intermediate and deep waters from the Pacific to the Caribbean along the CAS (Sepulcre et al. 2014; Montes et al. 2015; Jaramillo 2018). The temporal separation of transisthmian sister species among snooks (∼6.2 Ma), aligns with the distribution of vicariance events linked to the Isthmus of Panama. Bacon et al. (2015) proposed the events of divergence separating marine organisms to intensify during the to 10 to 4.2 Ma interval. O’Dea et al. (2016) report a divergence timeframe ranging from 10 Ma to 5 Ma for sister taxa within Teleostei. Additionally, various families of marine fishes, including Serranidae (Craig et al. 2004), Haemulidae (Tavera et al. 2012), Labridae and Chaetodontidae (Cowman and Bellwood 2013), Eleotridae and Apogonidae (Thacker 2017), exhibit a predominant vicariance pattern during the Late Miocene to Early Pliocene.

An allopatric event of speciation occurred ∼5.51 Ma in the snooks *Centropomus irae* and *C. undecimalis*; some authors suggest that the divergence in these species was induced by the influence of the Amazon River (Malcher et al. 2023). We believe that other mechanisms must be involved in the separation of this pair of species. *Centropomus undecimalis* is a widely distributed species in the Caribbean and the Atlantic, and it has a high tolerance to salinity (e.g. it has been reported from Lake Gatún in Panama). Rather, the separation of these species may be due to environmental and ecological factors. Even more recent speciation events within the family Centropomidae occurred within the same basin as in the sympatric Caribbean species *C. mexicanus* and *C. parallelus* (∼1.8 Ma), and the Pacific *C. armatus* and *C. robalito* (∼210 Ka).

Thus, speciation events in Centropomidae are found pre- and post-final closure of the Isthmus of Panama. This pattern aligns with observations in other studies focusing on diverse families of marine fishes, where divergence times have been detected before and after the uplift of the Isthmus of Panama. Some examples include the genus *Holacanthus* within the angelfish family Pomacanthidae (Tariel et al. 2016), the genus *Cyclopsetta* within the flatfish family Paralichthyidae (Byrne et al. 2018), and the genus *Haemulon* in the grunt family Haemulidae (Tavera et al. 2019). In the same way, our results are congruent with those found in diverse groups of marine invertebrates (Lessios 1981; Weinberg and Starczak 1989; Knowlton and Weight 1998; Marko and Jackson 2001; Lessios 2008; Miura et al. 2010; Hiller and Lessios 2019; Lima et al. 2020) where, after the closure of the Isthmus of Panama, climatic, oceanographic, and ecological conditions play important roles in sympatric speciation.

## Supplementary material

### Fossil occurrences

**Specie:** Centropomus sp.

### Material

United States, Florida. Polk country, Tiger Bay Mine, Hawthorn group, Peace River formation, Bone Valley member, early Pliocene collected by Eric Kendrew, articular left UF 293245 (Fig 1-A-C), basioccipital and parasphenoid UF 293352(Fig 2-A-D). Collected by Timberlane Research Organization Crew Ceratohyal left UF/TRO 23210 (Fig 3-A-B), vomer UF/TRO 23211(Fig 3-C-D); collected by John S. Waldrop Parasphenoid UF 291313(Fig 1-D-F). Nichols Mine, Hawthorn group, Peace River formation, Bone Valley member, early Pliocene collected by Eric Kendrew Dentary right UF 113175 (Fig 4 A-C), Hyomandibular, left UF 113209 (Fig 4 D-E), Premaxilla right UF 109998 (Fig 4 F-G) Palmetto Mine, Hawthorn group, Peace River formation, Bone Valley member Early Pliocene collected by John S. Waldrop Hypural UF 291601, vertebra atlas UF 293427, vertebra UF/TRO 23027 (Fig 5 A-D), UF 291382 (Fig 5 E-I). They were photographed by Sean Moran.

**(Figure 1 A-F).**
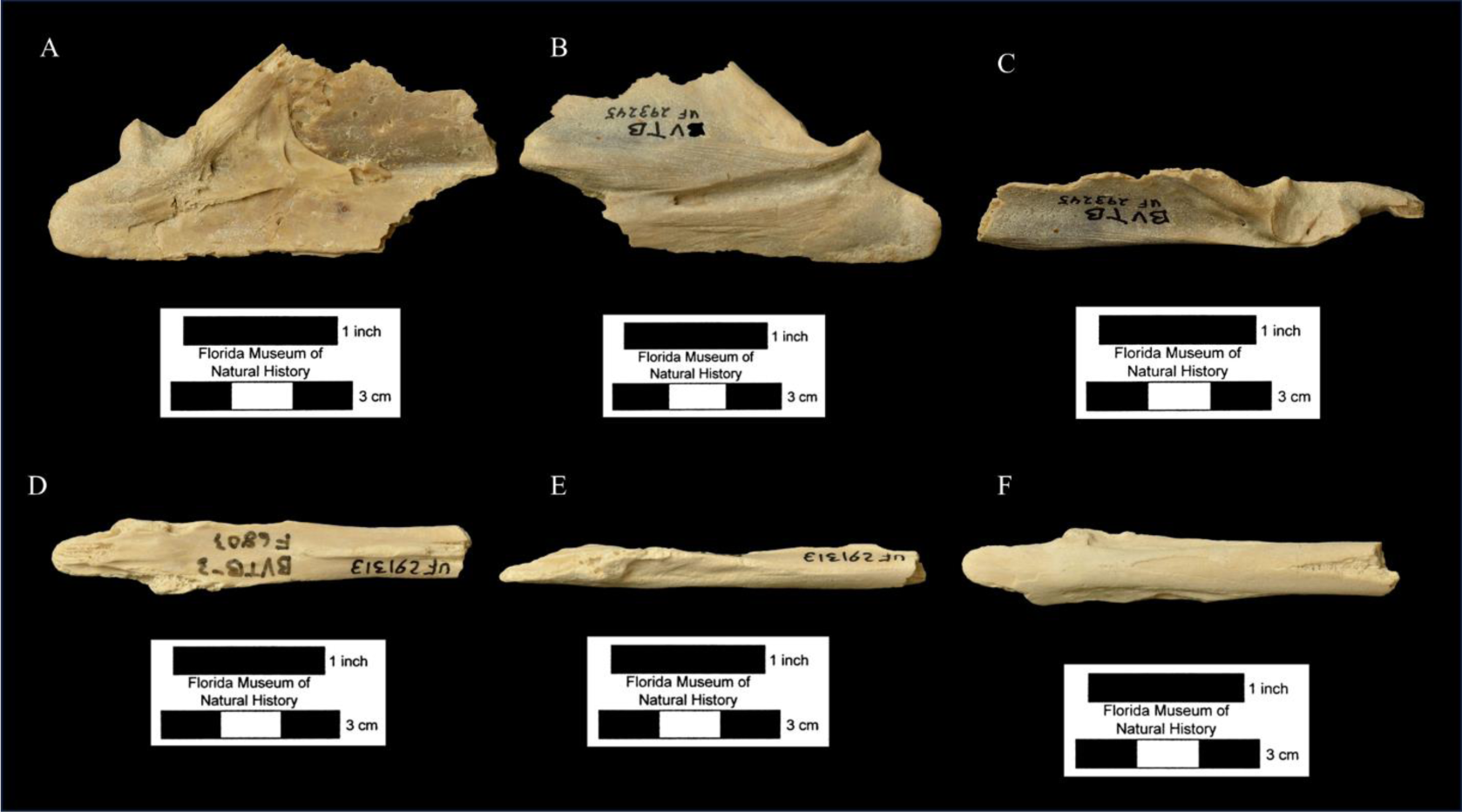

**(Figure 2 A-E).**
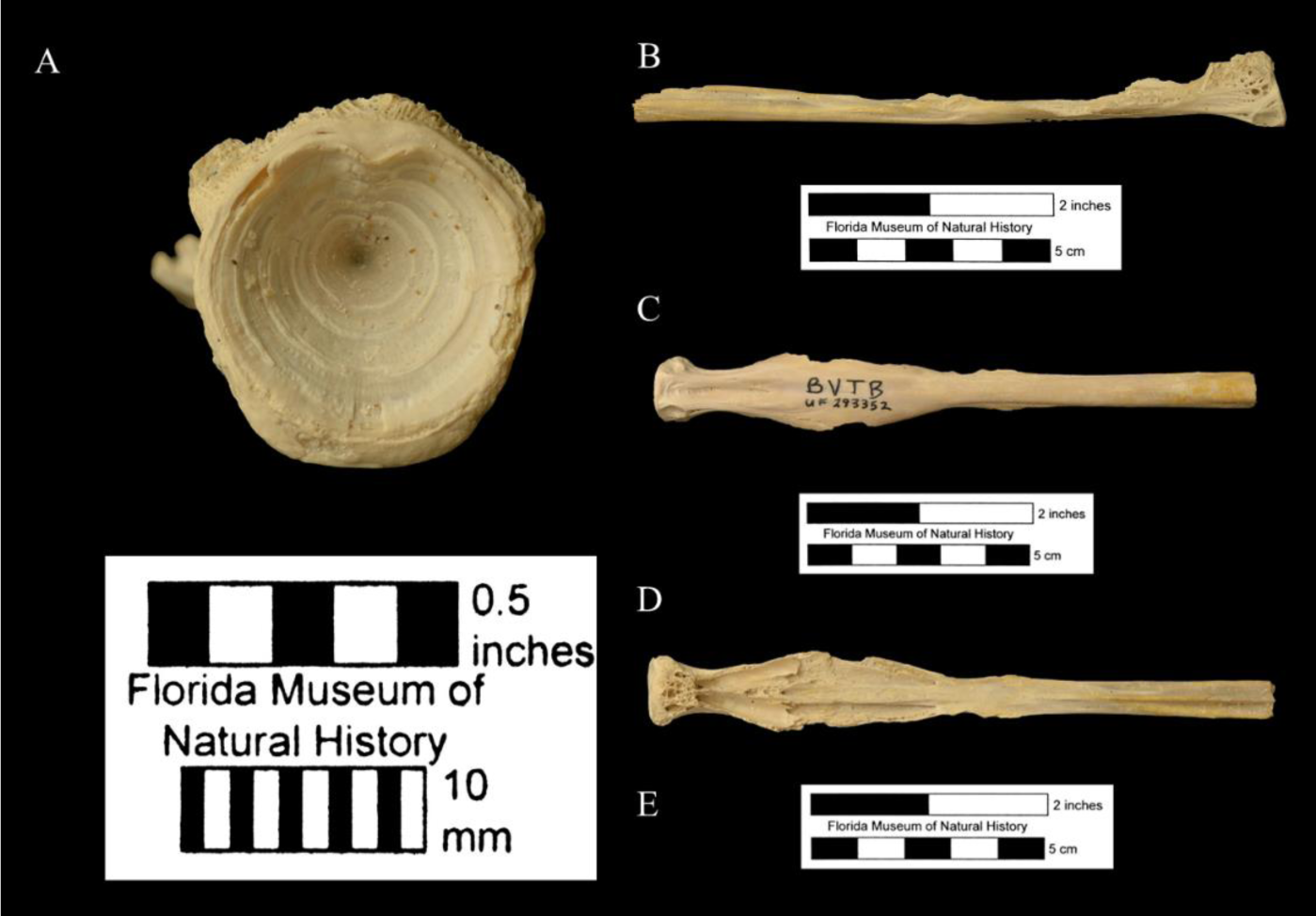

**(Figure 3 A-D).**
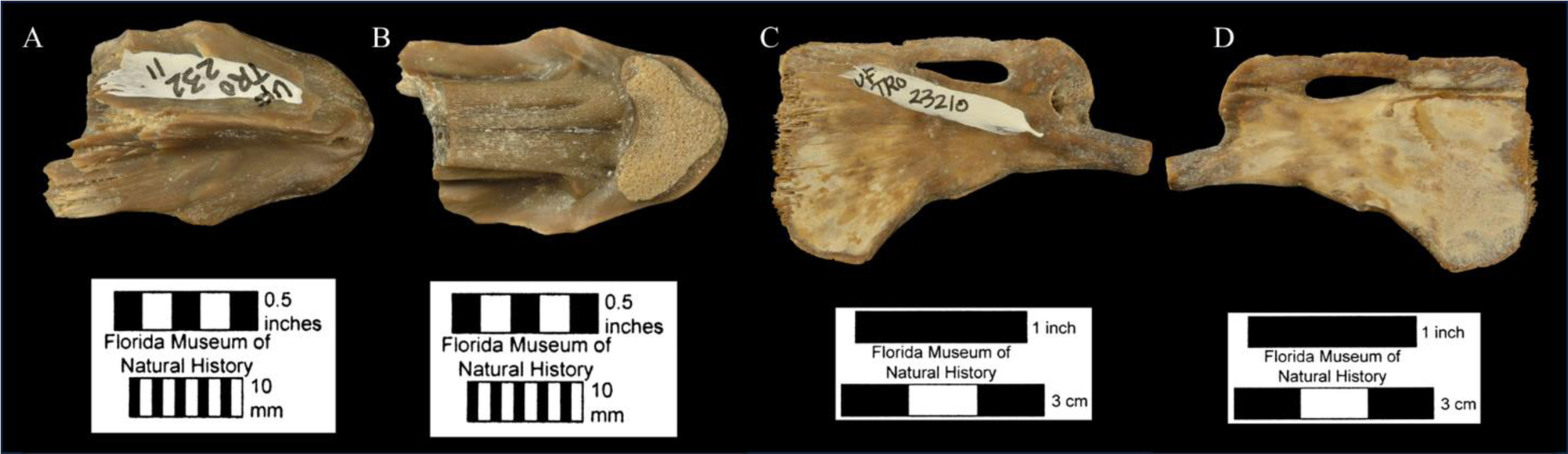

**(Figure 4 A-G).**
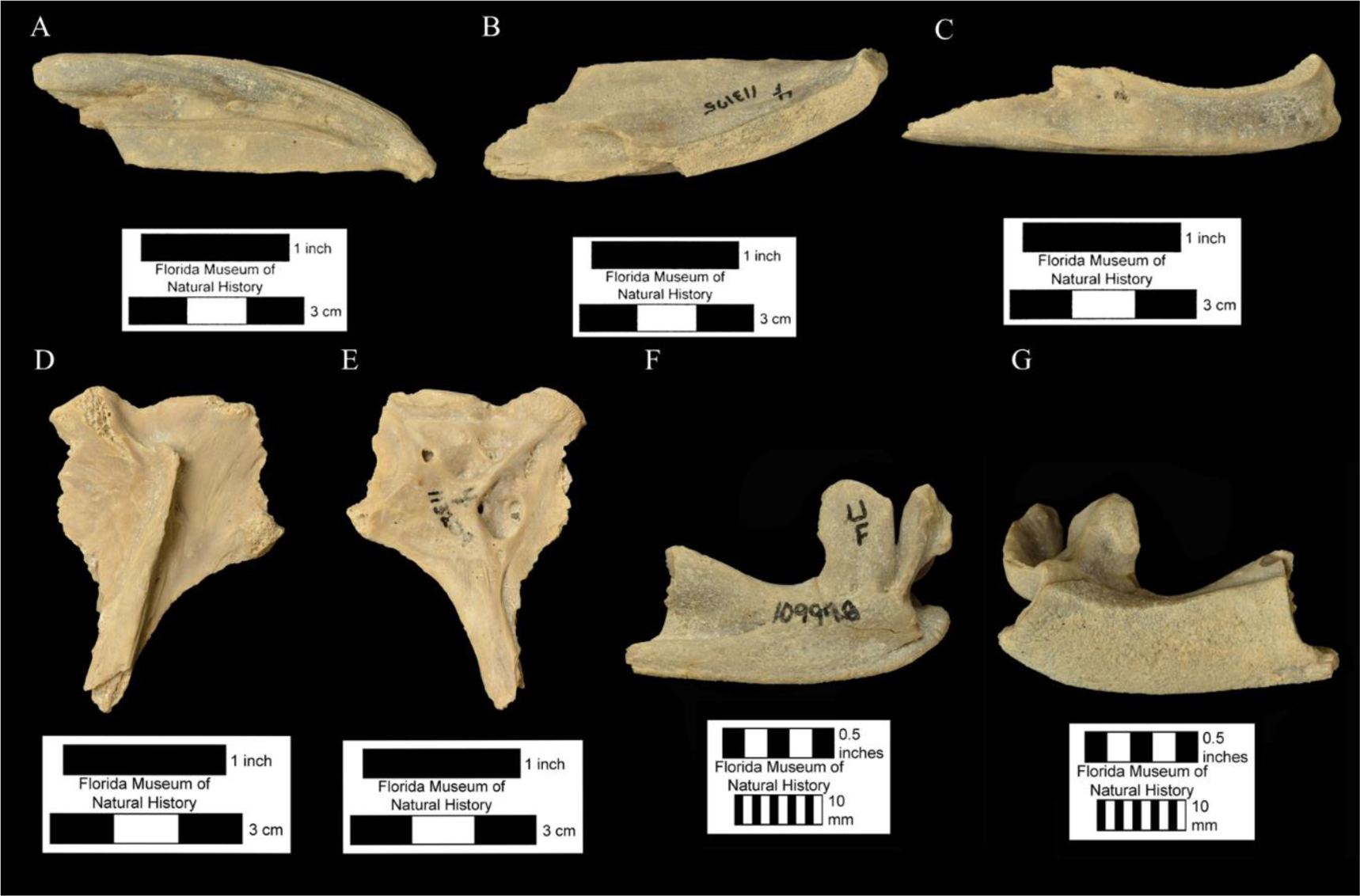

**(Figure 5 A-I).**
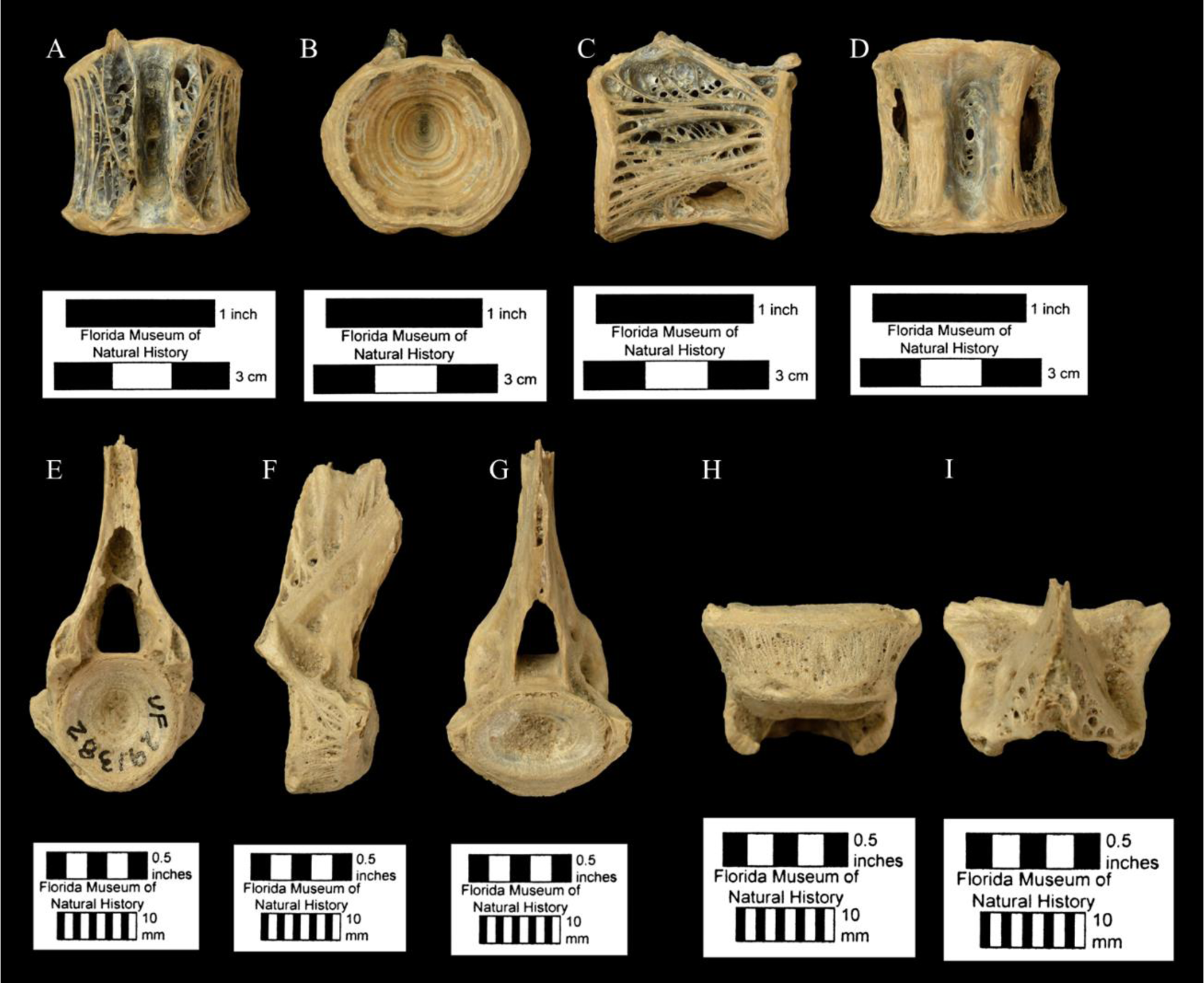

**Table S1.** Accession number in NCBI. *Sequences obtained in this study; **Sequences obtained in collaboration with the Ecological and Evolutionary Genomics Laboratory at STRI.

| *Especies* | *Localidad* | **12S** | **16S** | **COI** | **CYTB** | **TMO** |
| --- | --- | --- | --- | --- | --- | --- |
| *Centropomus armatus* | Colombia: Valle del cauca | OR805158* | OR805147* | OR825518* | OR829270* | OR8229617* |
| *Centropomus ensiferus* | Cabo de la Vela | OR805159* | OR805148* | OR825519* | OR829271* | OR8229618* |
| *Centropomus irae* |  |  | KJ641473.1 | MW183548 | KJ641490.1 |  |
| *Centropomus medius* | Colombia: Valle del cauca | OR805160* | OR805149* | OR825520* | OR829272* | OR8229619* |
| *Centropomus mexicanus* |  |  | U85015.3 | MZ376960 | AF018596.1 |  |
| *Centropomus nigrescens* | Panamá: Golfo de Panama | STRI** | STRI** | STRI** | OR829273* | OR8229620* |
| *Centropomus paralellus* |  |  | U85014.3 | JQ365275.1 | EU927346.1 |  |
| *Centropomus pectinatus* | Panamá: Colon | STRI** | STRI** | STRI** | OR829274* | OR8229621* |
| *Centropomus poeyi* |  |  | U85009.3 |  | AF018599.1 |  |
| *Centropomus robalito* | Colombia: Choco | OR805161* | OR805150* | OR825521* | OR829275* | OR8229622* |
| *Centropomus undecimalis* | Cabo de la Vela | OR805162* | OR805151* | OR825522* | OR829276* | OR8229623* |
| *Centropomus unionensis* | Colombia: Valle del cauca | OR805163* | OR805152* | OR825523* |  | OR8229624* |
| *Centropomus viridis* | Panamá: Golfo de Panama | STRI** | STRI** | STRI** | OR829277* |  |
| *Hypopterus macropterus* |  |  |  | LC269834. 1 |  |  |
| *Lates angustifrons* | Lake Tanganyika | MN255593.1 | MN255675.1 | ISZA08021 |  |  |
| *Lates calcarifer* |  | AY141371 | AY141441 | JF919821 | EU126588.1 |  |
| *Lates japonicus* |  | AP006788.1 | AP006788.1 | AP006788.1 | AP006788.1 |  |
| *Lates niloticus* |  | MN255585 | GU324156 | KJ443712 | AB117106.1 |  |
| *Lates stappersii* |  | MN255586.1 | MN255679.1 |  |  |  |
| *Psammoperca waigiensis* |  | KM082972.1 | KM198901.1 | FJ237578.1 | Y986971.1 |  |
| *Sphyraena acutipinnis* |  |  |  | HM902634.1 |  |  |
| *Sphyraena africana* |  | LC499574.1 |  |  |  |  |
| *Sphyraena argentea* |  |  | EU0099477.1 | GU440525 |  | DQ388071 |
| *Sphyraena barracuda* | Colombia: Choco | OR805164* | OR805153* | OR825524* | OR829278* | OR8229625* |
| *Sphyraena borealis* | Venezuela: Isla Margarita | OR805165* | OR805154* | OR825525* |  | OR8229626* |
| *Sphyraena crysotaenia* |  |  | GQ485295.1 | KY176643.1 |  |  |
| *Sphyraena ensis* | Colombia: Valle del cauca | OR805166* | OR805155* | OR825526* |  | OR8229627* |
| *Sphyraena flavicauda* |  |  | GQ485296.1 | KY176644.1 |  |  |
| *Sphyraena forsteri* |  | LC385307.1 |  | MK777495. 1 | AB264366.1 |  |
| *Sphyraena guachancho* | Colombia: Magdalena | OR805167* | OR805156* | OR825527* | OR829279* | OR8229628* |
| *Sphyraena helleri* |  | LC327187.1 |  |  | AB264367.1 |  |
| *Sphyraena iburrensis* |  | LC499397.1 |  |  | AB264361.1 |  |
| *Sphyraena idiaestes* |  |  | DQ532963.1 |  |  |  |
| *Sphyraena japonica* |  | AP012501.1. | AP012501.1. | AP012501.1. | AP012501.1. |  |
| *Sphyraena jello* |  | KT445895.1. | KT445895.1. | KT445895.1. | KT445895.1. |  |
| *Sphyraena novaehollandiae* |  |  | KX234684.1 |  |  |  |
| *Sphyraena obtusata* |  | LC037054.1 | KR153518.1 | KU499681.1 | AB264338.1. |  |
| *Sphyraena pinguis* |  | LC506671.1 |  | JF952863.1 | AB264356.1 |  |
| *Sphyraena putnamae* |  | LC037053.1 | JQ938993.1. | MZ329572.1 | KR007749.1 |  |
| *Sphyraena qenie* | Colombia: Valle del cauca | OR805168* | OR805157* | OR825528* | OR829280* |  |
| *Sphyraena sphyraena* |  | DQ533304.1 | JQ939013.1. | KY176645.1 | EF439597.1 |  |
| *Sphyraena viridensis* |  | KJ499108.1 |  | KJ396640.1 | DQ198006.1 |  |
| *Sphyraena waitii* |  |  |  | HM902624.1 |  |  |
| **Outgroup** |  |  |  |  |  |  |
| *Caranx caninus* | Colombia: Valle del cauca | This study | This study | OR825517* | This study |  |
| *Coryphaena equiselis* |  | MH576916.1 | MH576916.1 | MH576916.1 | MH576916.1 |  |
| *Echeneis naucrates* |  | KF021242. 1 | KF021242. 1 | KF021242. 1 | KF021242. 1 |  |
| *Gnathanodon speciosus* |  | NC_054367 | NC_054367 | NC_054367 | NC_054367 |  |
| *Isthiophorus platypterus* |  | NC_012676.1 | NC_012676.1 | NC_012676.1 | NC_012676.1 | DQ3880772.1 |
| *Makaira nigricans* |  | HQ611116. 1 | HQ592244.1 | HQ611116. 1 | HQ611116.1 | DQ388073.1 |
| *Nematistius pectoralis* |  | ON838225.1 | ON838225.1 | ON838225.1 | ON838225.1 |  |
| *Tetrapterus audax* |  | NC_012678.1 | NC_012678.1 | NC_012678.1 | NC_012678.1 |  |
| *Xiphias gladius* |  | AB470301 | AB470301.1 | AB470301 | AB470301 | DQ388074.1 |

**Table S2.**
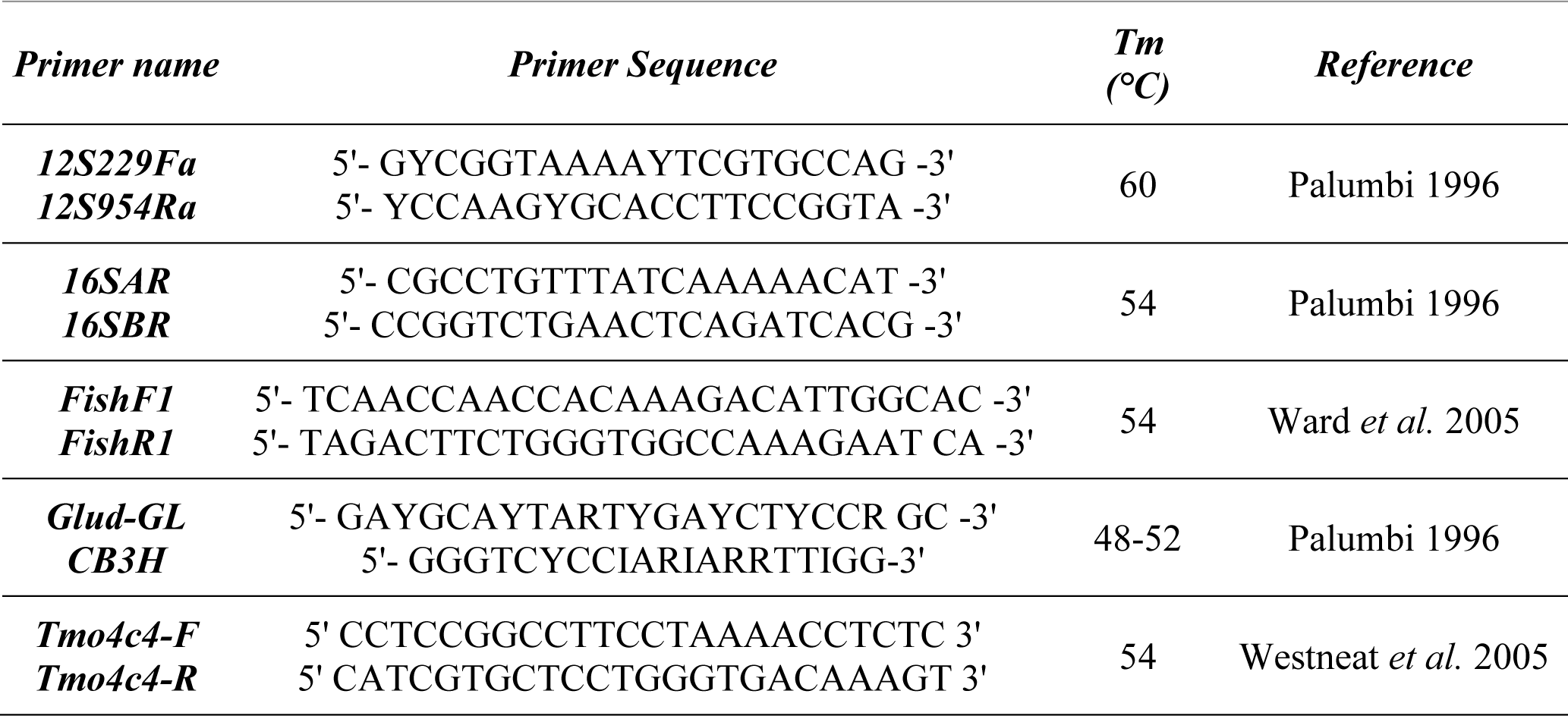
Primer sequences used for PCR amplifying this study’s mitochondrial and nuclear genes.

**Table S3.** Bayesian relaxed clock model selection using marginal likelihood and Bayes factors. The relaxed clock where independent rates follow a lognormal distribution was strongly favored as the best for the current dataset.

| Clock Model | Log (Marginal Likelihood) | Standard Error | Posterior Probability |
| --- | --- | --- | --- |
| Independent lognormal | -536.573980786701 | 0.475257291697664 | 0.999999999993294 |
| Geometric Brownian Motion | -562.302019175086 | 0.551420967186908 | 0.00000000000670586641 |

**Table S4.**
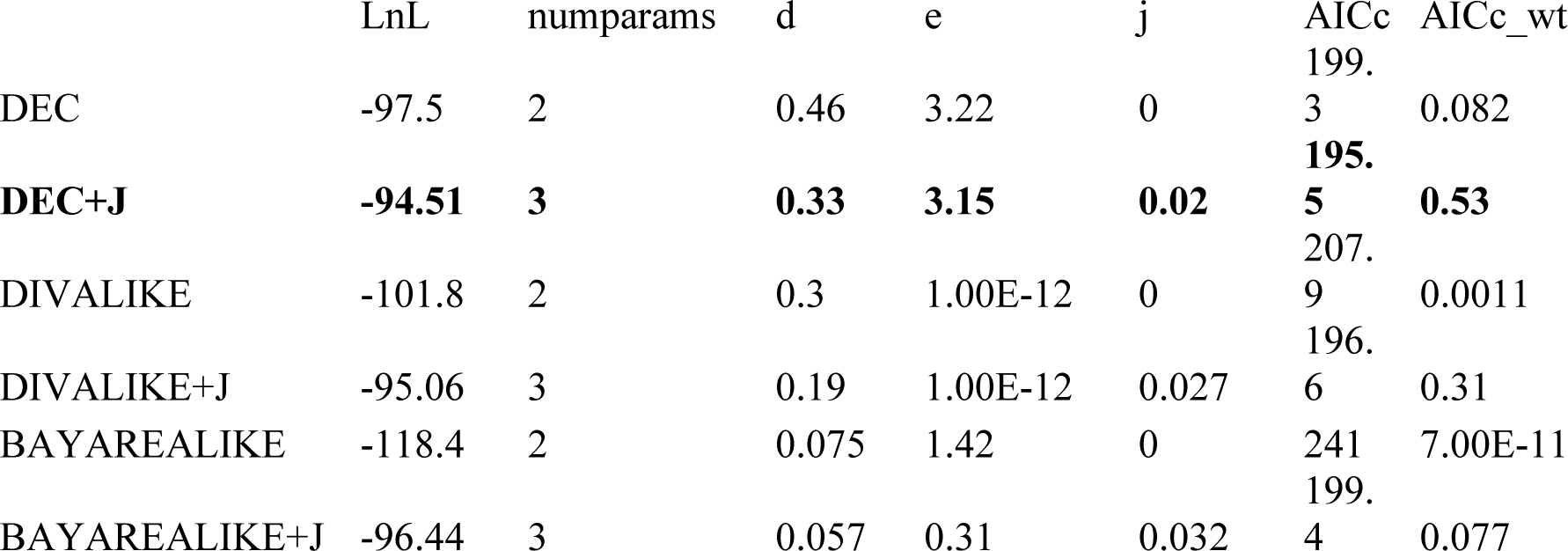
Comparison of four biogeographic model selections using the weight of the Akaike Information Criterion (AICwt). The best model is highlighted in boldface.

## Conclusions

The family Centropomidae is the sister clade of the family Sphyraenidae, with a common ancestor in the Upper Cretaceous. The families Latidae and Sphyraenidae originated in the Paleocene in the region we know today as the Indo-Pacific. The family Centropomidae originated in the Western Atlantic and Eastern Pacific in the Oligocene. The transitions in the estimated ancestral area in this last family occurred from the Eastern Pacific to the Western Atlantic during the emergence of the isthmus of Panama. The divergence times and biogeographic patterns identified in the family Centropomidae are congruent with a gradual impact on the species diversification, rather than a single synchronous event during the formation process of the Isthmus of Panama. Both geographic isolation during and after the emergence of the Isthmus of Panama and the environmental and ecological changes created post-Isthmus of Panama shaped the diversification in this family.

## Acknowledgments

NOH was funded by Corporación CE-Marin through Call 14, 2018 supporting PhD candidates. The project was financed by the Universidad de Valle (an internal call for the presentation of research projects in 2018, and to support doctoral students in 2021). GAB was funded by the Fundação de Amparo à Pesquisa do Estado de São Paulo (processes 2014/11558-5, 2016/02253-1, and 2023/07838-1). All the analyses were run on the *Brycon* server at IBB/UNESP funded by FAPESP proc. 2014/26508-3. Thanks to Dra. Kristin Saltonstall and Dr. D. Ross Robertson for support and guidance in obtaining sequences at Smithsonian Tropical Research. Thanks to Rachel E. Narducci, Division of Vertebrate Paleontology Florida Museum, and Emma Bernard, the Natural History Museum, United Kingdom for their help with the revision of the fossil material in these collections. Contribution No. 569, Instituto para el Estudio de las Ciencias del Mar (Cecimar), Universidad Nacional de Colombia sede Caribe. Carlos Jaramillo is especially acknowledged for having hosted NOH during a predoctoral internship at STRI Panamá, where this project could be completed.

